# Blood-based Transcriptomics Reveal Sex- and Amyloid-Modulated Biology of Plasma pTau217 in Preclinical Alzheimer’s Disease

**DOI:** 10.1101/2025.11.21.689770

**Authors:** Mabel Seto, Hannah M. Klinger, Michelle Clifton, Vaibhav A Janve, Jane A. Brown, Colin Birkenbihl, Gillian Coughlan, Diana L. Townsend, Ting-Chen Wang, Michael Properzi, Bernard Hanseeuw, Jasmeer Chhatwal, Hyun-Sik Yang, Yanling Wang, Philip L. de Jager, Lisa L. Barnes, Julie A. Schneider, David A. Bennett, Robert A. Rissman, Paul Aisen, Madison Cuppels, Michael C. Donohue, Keith A. Johnson, Reisa A. Sperling, Logan Dumitrescu, Timothy J. Hohman, Rachel F. Buckley

**Affiliations:** Massachusetts General Hospital/Harvard Medical School, Boston, MA, USA; Brigham and Women’s Hospital, Boston, MA, USA; Vanderbilt Memory & Alzheimer’s Center, Vanderbilt Health, Nashville, TN, USA; Rush Alzheimer’s Disease Center, Rush University Medical Center, Chicago, IL, USA; Center for Translational & Computational Neuroimmunology, Columbia University Medical Center, New York, NY, USA; Alzheimer’s Therapeutic Research Institute, Keck School of Medicine of the University of Southern California, San Diego, CA, USA

## Abstract

Plasma pTau217, an emerging Alzheimer’s disease (AD) biomarker, may reflect a synaptic response to β-amyloid (Aβ) plaques before cortical tangle formation, but its broader biological correlates remain unclear. We sought to identify associations between whole blood gene expression and plasma pTau217, and to determine whether *APOE*ε4, sex, and neocortical Aβ-PET modify these associations in 724 participants from the Anti-Amyloid Treatment in Asymptomatic Alzheimer’s and accompanying LEARN studies (A4/LEARN, Age_mean(SD)_=72.2(4.6); 63%female). Of 20,621 genes tested (1,048 X-linked), none were directly associated with pTau217; one gene was moderated by *APOE*ε4, 1,540 genes by Aβ-PET, and 772 genes by both Aβ-PET and sex. Over 100 of these significant associations were X-linked, supporting a role of the X chromosome in AD. Sex interactions were only observed in the presence of elevated Aβ-PET. Our results underscore the complexity of molecular mechanisms that can be linked to plasma pTau217, particularly in the context of elevated Aβ-PET.

## Main

Plasma concentration of phosphorylated tau at threonine 217 (pTau217) has emerged as a sensitive blood-based biomarker for Alzheimer’s disease (AD)^1–4^. It has outperformed other phospho-tau species (e.g., pTau181, pTau231) in detecting abnormal AD brain pathology at the earliest stages of disease progression and has demonstrable selectivity for AD pathology^5–8^. Specifically, plasma pTau217 shows strong associations with β-amyloid (Aβ)-PET, and to a lesser extent, tau-PET.^2, 9–12^ Elevated plasma pTau217 also predicts future cognitive decline^13^ and progression to mild cognitive impairment (MCI) and dementia^11, 14–16^, further underscoring its importance as an early marker of AD. Though plasma pTau217 shows clear promise as a minimally invasive AD biomarker, the biological pathways associated with early pTau217 elevation in the blood of cognitively unimpaired (CU) older adults remains unclear.

Neuropathological studies demonstrate that pTau217 species appear in synapses surrounding Aβ plaques even prior to overt tau neurofibrillary tangle formation, supporting it as a very early signal of Aβ deposition^17^. Postmortem AD brain measures of pTau217 have also been linked to signals of synaptic loss and stress (e.g., granulovacuolar and multivesicular bodies)^18^ in neuropathological studies of AD, perhaps in direct response to the aggregation of Aβ^18^. Given these findings, elevated plasma pTau217 may index a broader set of early Aβ-adjacent pathological processes, including inflammation, synaptic stress and degeneration, particularly in individuals with abnormal Aβ or who carry an *APOE*ε4 allele, which is the most common genetic risk factor for AD.

Sex represents another critical, and often overlooked, factor when considering biological mechanisms driving the earliest AD pathophysiological changes. Women not only show a higher prevalence of AD dementia but also appear more vulnerable to tau pathology and cognitive decline at higher levels of Aβ burden^19–21^. Women with elevated Aβ also show higher baseline plasma pTau217 than men with elevated Aβ, and women with higher plasma pTau217 show faster rates of tau-PET accumulation^22^, suggesting that sex may offer additional and important context for biological pathways driving elevated plasma pTau217 and other early pathophysiological changes in AD. Beyond hormonal influences to explain these sex effects, the genome, including the X chromosome, is increasingly being found to play a role in sex-specific AD risk. A growing body of work shows that genetic associations with AD endophenotypes differ by sex^23^, suggesting the involvement of distinct sex-specific biological pathways^24^. The X chromosome, which accounts for approximately 5% of the genome^25^ and harbors many brain-expressed genes^26^, has been remarkably understudied in this space^24^. The X chromosome is increasingly implicated in AD through genetic studies^23, 24, 27^. Investigating peripheral transcriptional correlates of plasma pTau217, including X-linked signals, in a sex-specific manner may therefore provide insight into the biological context of early AD-related change in CU older adults.

The overall aim of this study was to identify associations between whole blood gene expression, on both the autosome and the X chromosome, and plasma pTau217 levels in more than 700 cognitively unimpaired older adults from the Anti-Amyloid Treatment in Asymptomatic Alzheimer’s (A4) clinical trial and the adjoining Longitudinal Evaluation of Amyloid Risk and Neurodegeneration (LEARN) observational study. We further explored moderating associations with Aβ-PET, *APOE*ε4, and sex to better characterize the biological contexts in which plasma pTau217 is elevated during the preclinical phase of AD.

## Results

Participant demographics are shown in **Table 1** and stratified by study (i.e., A4 and LEARN). For this sample of 724 individuals (n_A4_=445, n_LEARN_=279), the mean age was 72.2 years (SD±4.6; range: 65-86 years) and mean education was 16.5 years (SD±2.6; range: 7-30 years); 63% were female. As per eligibility criteria, individuals enrolled in A4 exhibited abnormal levels of Aβ-PET (Aβ-PET standardized uptake value ratio, SUVR ≥ 1.15 and qualitative visual read). Individuals with an SUVR between 1.10 and 1.15 were considered to have elevated brain Aβ-PET when a visual read was considered positive by a 2-person consensus^28^. Individuals who did not have elevated Aβ-PET but were otherwise eligible for the A4 study were invited to enroll in the LEARN companion study. Due to these enrollment criteria, 62% of individuals in our sample were Aβ-positive (SUVR > 1.10) and 45% carried at least one *APOE*ε4 allele. There were no significant differences in mean age or years of education between individuals enrolled in A4 compared to LEARN, though baseline pTau217, global Aβ-PET burden, and *APOE*ε4 status (i.e., positive or negative) statistically differed between A4 and LEARN (p<0.001).

**Table 1:** Participant Demographics.

| Table 1: Participant Demographics |  |  |  |  |
| --- | --- | --- | --- | --- |
| Characteristic | Overall<br>N = 724<sup>1</sup> | A4 (Aβ+)<br>N = 445<sup>1</sup> | LEARN (Aβ-)<br>N = 279<sup>1</sup> | p-value<sup>2</sup> |
| Baseline Age | 72.15 (4.64)<br>[65.21, 85.99] | 72.24 (4.83)<br>[65.21, 85.75] | 72.00 (4.32)<br>[66.54, 85.99] | 0.8 |
| Sex [Female] | 457 (63.12%) | 284 (63.82%) | 173 (62.01%) | 0.6 |
| Years of Education | 16.53 (2.60)<br>[7.00, 30.00] | 16.53 (2.71)<br>[7.00, 30.00] | 16.53 (2.42)<br>[9.00, 26.00] | >0.9 |
| *APOE*ε4 Status [ε4+] | 325 (44.95%) | 258 (57.98%) | 67 (24.10%) | **<0.001** |
| Baseline pTau217 | 0.24 (0.15)<br>[0.13, 1.43] | 0.28 (0.17)<br>[0.13, 1.43] | 0.16 (0.06)<br>[0.13, 0.58] | **<0.001** |
| Baseline Global Aβ-PET SUVR | 1.20 (0.21)<br>[0.83, 2.00] | 1.33 (0.16)<br>[1.01, 2.00] | 1.00 (0.07)<br>[0.83, 1.15] | **<0.001** |

<sup>1</sup>Mean (SD) [Range], n (%); <sup>2</sup>Wilcoxon rank sum test; Pearson's Chi-squared test; <sup>3</sup>LEARN participants are not randomized into treatment groups.

In primary analyses, we used linear regression models to examine associations between whole-blood gene expression (n_total_ = 20,621 genes; 19,573 autosomal and 1,048 X-linked) and cross-sectional plasma pTau217 levels, as well as potential moderation by *APOE*ε4 status, global Aβ-PET SUVR, and sex. Specifically, models tested the main effect of gene expression on plasma pTau217, two-way interactions between gene expression and Aβ-PET SUVR, *APOE*ε4 status, or sex, and a three-way interaction between gene expression, Aβ-PET SUVR, and sex, with all models adjusted for covariates. Covariates included baseline age, body mass index (BMI; to account for blood volume), and cohort (i.e., A4, LEARN). All primary analyses, including gene set enrichment analyses, were corrected for multiple comparisons using the false discovery rate (FDR) method; p_FDR_ < 0.05 was considered significant. *Post hoc* analyses (e.g., stratified models and additional exploratory analyses) were not corrected for multiple comparisons unless otherwise stated.

### Gene associations with pTau217 as moderated by Aβ-PET

No autosomal or X-linked genes were directly associated with pTau217 levels. 1,540 genes in total interacted with Aβ-PET SUVR on pTau217 (120 X-linked genes (7.8%), **Supplementary Table 1**). 63% of these genes were found to be protective; that is, higher gene expression was associated with lower plasma pTau217 among individuals with elevated Aβ-PET burden. Six of the 1,540 significant genes (*EED, IL34, EPHA1, WDR81, TREML2, CASP7*) were noted to have “significant evidence of affecting AD risk,” by the Alzheimer’s Disease Sequencing Project (ADSP) Gene Verification Committee^29–34^. Of these genes, *EPHA1* and *TREML2* exacerbated the Aβ association with pTau217 whereas *EED, IL34, WDR81*, and *CASP7* demonstrated a protective effect against Aβ-moderated pTau217 levels (**Table 2**).

**Table 2.** Gene x Aβ-PET Association Results for Genes Appearing on the ADSP List of AD Loci and Genes with Genetic Evidence.

| **Gene Name** | **Chr** | **Beta** | **SE** | **P** | **P<sub>FDR</sub>** |
| --- | --- | --- | --- | --- | --- |
| *TREML2* | 6 | 0.237 | 0.067 | <0.001 | 0.015 |
| *EPHA1* | 7 | 0.252 | 0.066 | <0.001 | 0.009 |
| *CASP7* | 10 | -0.233 | 0.077 | <0.001 | 0.041 |
| *EED* | 11 | -0.421 | 0.101 | <0.001 | 0.003 |
| *IL34* | 16 | -0.078 | 0.019 | <0.001 | 0.005 |
| *WDR81* | 17 | -0.498 | 0.132 | <0.001 | 0.009 |

Among novel autosomal genes, Small Integral Membrane Protein-38 *(SMIM38)* showed the largest negative Aβ-moderated association with pTau217 (β=-0.13(0.02), p_FDR_=2.84×10^−5^), while *FAM13A-AS1* showed the largest positive association (β=0.67(0.13), p_FDR_=1.62×10^−4^). *SMIM38* expression was associated with lower plasma pTau217 levels among individuals with elevated Aβ-PET (**Figure 1A**), while *FAM13A-AS1* was associated with higher plasma pTau217 levels in individuals with elevated Aβ-PET (**Figure 1B**). *SMIM38* has no known function, though is localized in the cell membrane. *FAM13A-AS1* is an antisense RNA that corresponds to the *FAM13A* gene and has been related to adipocyte function^35^.

**Figure 1.**
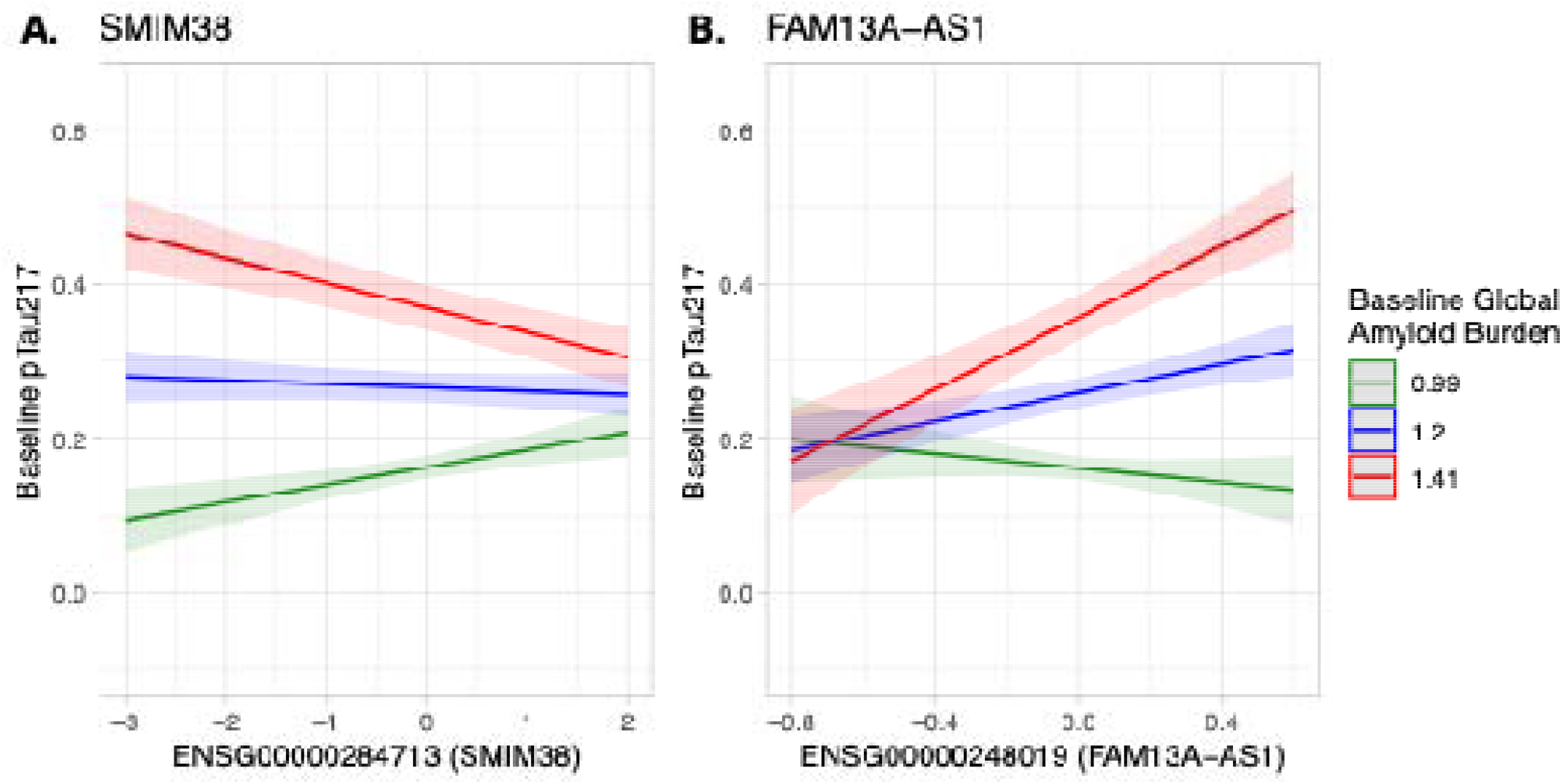
Model estimates predicting pTau217 levels from (A) an *SMIM38*×Aβ interaction and (B) a *FAM13A-AS1*×Aβ interaction. The mean ± 1 standard deviation (SD) of Aβ-PET SUVR is given such that blue represents the mean Aβ-PET SUVR, red is 1SD above the mean, and green is 1SD below the mean.

The most significant X-linked gene in the gene×Aβ-PET interaction analyses was *MIRLET7F2* (β=-0.09(0.01), p_FDR_=2.84×10^−5^, **Figure 2A**). Higher *MIRLET7F2* (microRNA) was associated with lower pTau217 levels among individuals with higher Aβ-PET burden. We found two X-linked genes that were previously implicated in AD studies that also showed significance: *FAM156B*^*27*^ and *KDM6A*^*36*^. Higher *FAM156B* expression (β=-0.09(0.03), p_FDR_<0.001, **Figure 2B**) was associated with lower pTau217 levels among individuals with higher Aβ-PET burden whereas higher *KDM6A* expression (β=0.31(0.10), p_FDR_=0.003, **Figure 2C**) was associated with higher pTau217 levels. These three genes, two of which are known to be highly expressed in brain, highlight transcriptional regulation and histone modification^36^ as biological pathways associated with plasma pTau217.

**Figure 2.**
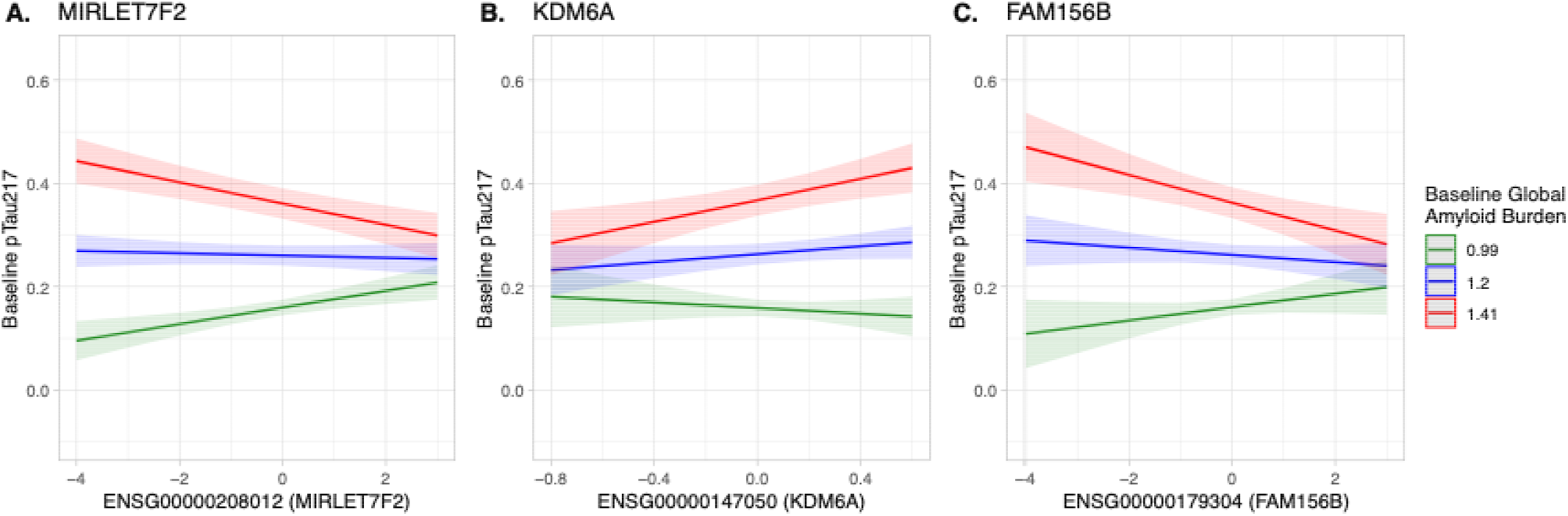
Model estimates predicting pTau217 levels from (A) an *MIRLET7F2*×Aβ interaction, (B) a *KDM6A*×Aβ interaction, and (C) a *FAM156B*×Aβ. The mean ± 1 standard deviation (SD) of Aβ-PET SUVR is given such that blue represents the mean Aβ-PET SUVR, red is 1SD above the mean, and green is 1SD below the mean.

To link our significant findings with broader biological processes, we performed gene set enrichment analyses using Gene Ontology: Biological Process (GO:BP) terms^37, 38^. Our enrichment analyses suggested a downregulation of functions such as aerobic respiration and fatty acid metabolism, though no terms survived correction for multiple comparisons (**Supplementary Table 2**). As a sensitivity analysis, we stratified by cohort (i.e., A4 and LEARN). 1,438 of 1,540 genes remained significant in individuals from A4 who had elevated Aβ, but none were significant in the LEARN subgroup (**Supplementary Table 3**) supporting our results showing gene associations primarily moderated by elevated brain Aβ.

### Gene associations with pTau217 as moderated by APOEε4

One long, uncharacterized noncoding RNA (lncRNA; ENSG00000273139) showed a significant interaction with *APOE*ε4 status (β=-0.14(0.03), p_FDR_=0.03). We further stratified by *APOE*ε4 status and found that, among *APOE*ε4 carriers, greater expression of ENSG00000273139 was associated with lower plasma pTau217 levels (β=-0.11(0.02), p_uncorrected_=5.02×10^−6^).

### Gene associations with pTau217 as moderated by sex and Aβ-PET

There were no significant gene associations with pTau217 levels that were moderated by sex alone. 772 genes (27 X-linked genes (3.5%), **Supplementary Table 4**), however, showed a significant three-way interaction between gene expression, sex, and Aβ-PET SUVR (i.e., gene×Aβ-PET SUVR×sex) on plasma pTau217 levels. None of the genes identified in these models were highlighted by the ADSP Gene Verification Committee^29–34^. *AMIGO2* was the most significant autosomal gene (β=-0.93(0.14), p_FDR_=3.79×10^−6^). Greater *AMIGO2* expression corresponded with higher pTau217 levels in the setting of elevated Aβ-PET burden in females (β=0.49(0.10), p_FDR_=5.57×10^−4^), while the opposite was true in males (β=-0.44(0.10), p_FDR_=3.79×10^−3^; **Figure 3A**). The second-most significant gene was *CERKL* (β=0.58(0.10), p_FDR_=3.28×10^−5^). Among females with elevated Aβ-PET, greater *CERKL* expression was associated with lower pTau217 burden (β=-0.36(0.06), p_FDR_=3.17×10^−5^), while the opposite effect was observed in males (β=0.21(0.07), p_FDR_=0.048; **Figure 3B**). *AMIGO2* encodes a cell adhesion molecule involved in axon guidance and survival^39^, while *CERKL* is suggested to playba role in mitochondrial function and autophagy regulation^40^. Sex-stratified results for gene×Aβ-PET SUVR×sex models are in **Supplementary Table 5**.

**Figure 3.**
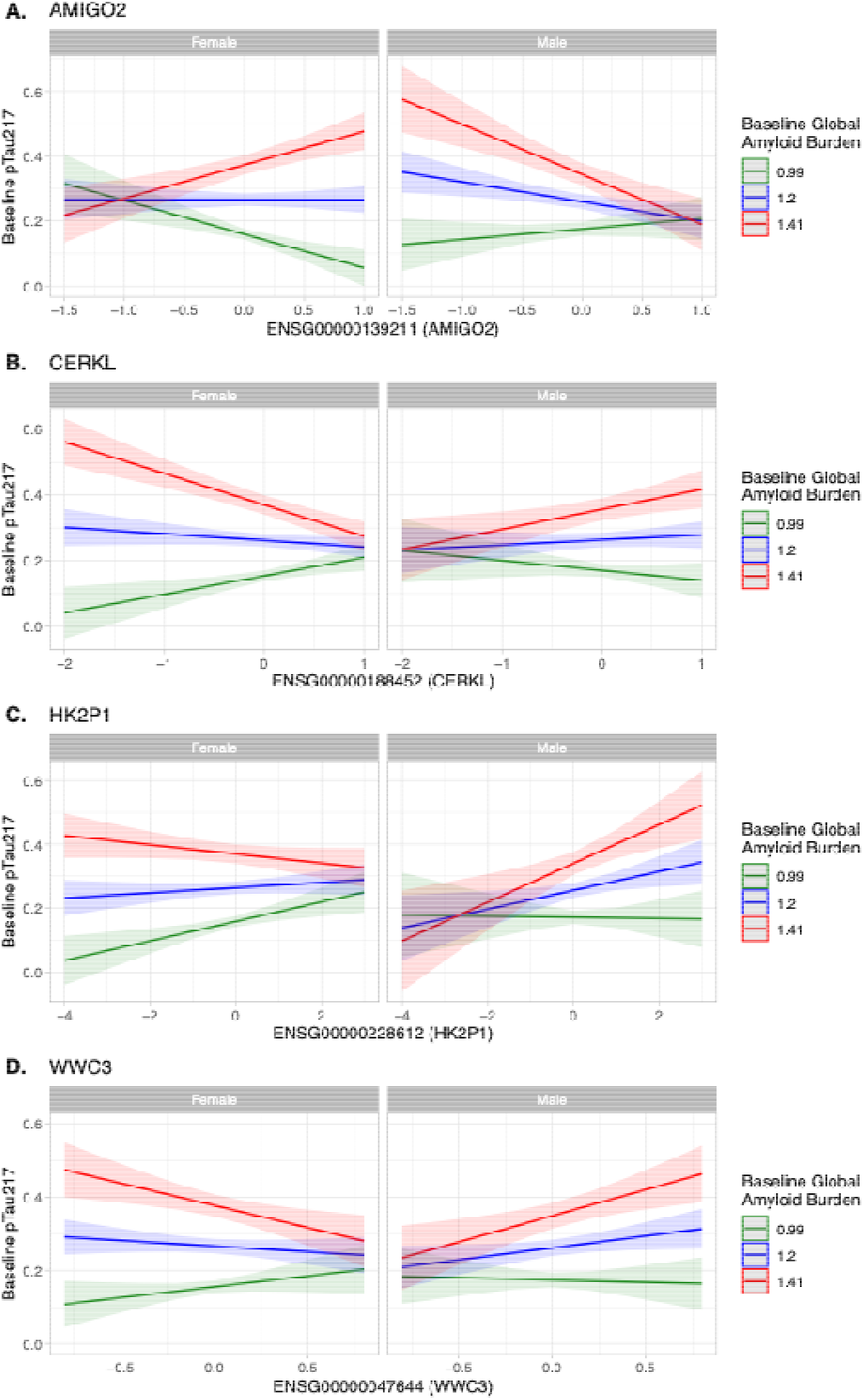
Model estimates predicting pTau217 levels from (A) an *AMIGO2*×Aβ×sex interaction, (B) a *CERKL*×Aβ×sex interaction, (C) a *HK2P1*×Aβ×sex interaction, and (D) a *WWC3*×Aβ×sex interaction. The mean ± 1 standard deviation (SD) of Aβ-PET SUVR is given such that blue represents the mean Aβ-PET SUVR, red is 1SD above the mean, and green is 1SD below the mean.

The most significant X-linked gene in our three-way interactions was *HK2P1* (β=0.25(0.06), p_FDR_=0.01), which is a pseudogene of *HK2. HK2* plays a role in glycolysis^41^ and *HK2P1* is suggested to play a similar function in addition to endometrial cellular preparation for pregnancy (decidualization)^42^. When stratifying by sex, greater expression of *HK2P1* was associated with lower pTau217 levels among females with elevated Aβ-PET (β=-0.11(0.03), p_uncorrected_=4.0×10^−4^) and greater pTau217 among males (β=0.15(0.05), p_uncorrected_=0.008) with elevated Aβ-PET (**Figure 3C**). Another X-linked gene of note was *WWC3* (β=0.79(0.21), p_FDR_=0.02, **Figure 3D**), which regulates the *Wnt* and *Hippo* signaling pathways^43^ and is suggested to play an immune-related role in AD via B lymphocytes^44^. In females with elevated Aβ-PET, greater *WWC3* expression was associated with lower pTau217 burden (β=-0.44(0.15), p_FDR_=0.004), whereas males with elevated Aβ-PET had higher levels of pTau217 (β=0.37(0.14), p_FDR_=0.01). Other highly significant X-linked genes included *SLC35A2* (cellular transport/glycosylation)^45^ and *PNMA6A* (apoptosis, associated with neurodevelopmental and metabolic disorders)^46^, altogether underscoring immune, metabolic, and post-translational modification mechanisms relevant to early AD. The most significant results from gene set enrichment analyses largely related to ribosome biogenesis RNA processing (**Supplementary Table 5**), which were downregulated.

In sensitivity analyses that were stratified by cohort (i.e., A4 and LEARN), 727 of 772 genes remained significant in individuals from A4 (n=445), but none were significant in the LEARN subgroup (n=229, **Supplementary Table 7**).

### Genetically Predicted Expression

As the presence of Aβ pathology itself could affect gene expression levels, we wanted to examine the extent to which our significant results were driven primarily by genotype. To do so, we calculated gene expression values using the MetaXcan method (previously known as PrediXcan)^47^. Briefly, MetaXcan models were built with gene expression data from the NIH Genotype Tissue Expression (GTEx) Project (version 8, build 38; NIH database of Genotypes and Phenotypes (dbGaP) accession number: phs000424.GTEx.v8.p2, downloaded 11/13/2019)^48^. The MetaXcan method utilizes an elastic-net supervised machine learning model to predict gene expression directly from quality-controlled genotype data. Predicted gene expression from whole blood was available for 637 out of 724 individuals from our sample.

There were 2,312 significant associations in total for our primary gene×Aβ-PET and gene×Aβ-PET×sex analyses, of which 2,275 (98.4%) were considered unique genes. We were able to obtain predicted gene expression values via MetaXcan for 590 of 2,275 unique genes. 397 (67.3%) of those were originally significant in gene×Aβ-PET interaction analyses, while the rest were from gene×Aβ-PET×sex interaction analyses (**Supplementary Table 8**). No X-linked genes were predicted via MetaXcan. For these *post hoc* analyses, we used the same linear regression models previously reported; models were corrected for multiple comparisons using the FDR method. 33 genes showed significance (p_FDR_ < 0.05) in gene×Aβ-PET analyses and had beta coefficients that remained consistent (i.e., negative betas remained negative, etc). One gene of particular interest was *SMAD3* (chromosome 15, β=-0.42(0.08), p_FDR_ = 2.92×10^−4^, **Figure 4A, B**), due to previous studies relating it to Aβ clearance^49, 50^. In our three-way interaction analyses, 14 genes showed significance and had consistent beta coefficients (**Supplementary Table 9**). Another gene of particular interest was *TRIM65* (chromosome 17, β=-0.57(0.12), p_FDR_=6.67×10^−4^, **Figure 4C, D**), which encodes a ubiquitin ligase and plays a role in innate immunity^51^. None of these significant genes were previously highlighted by the ADSP Gene Verification Committee^29–34^.

**Figure 4.**
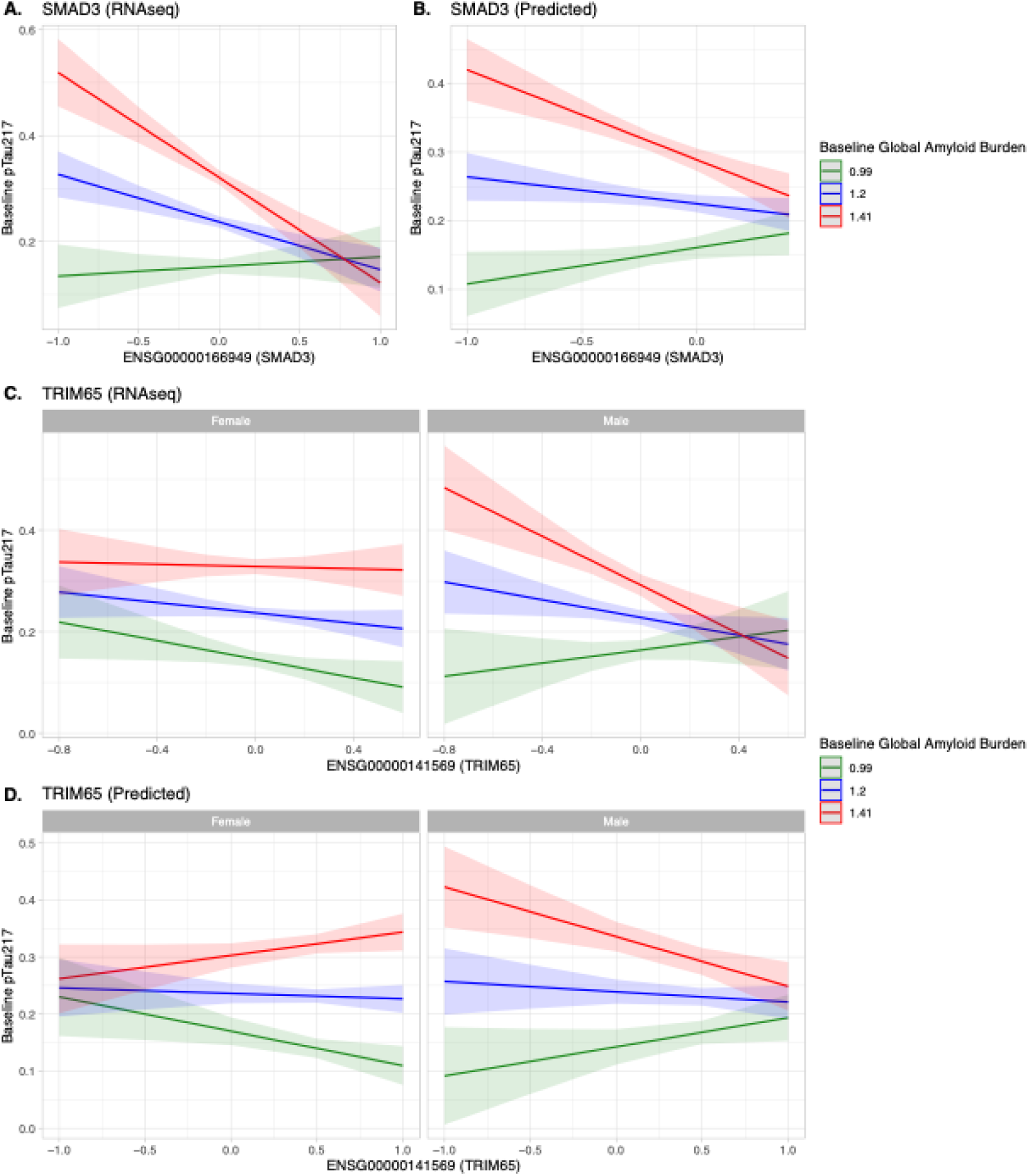
Model estimates predicting pTau217 from an (A) *SMAD3*×Aβ interaction using RNAseq data and (B) *SMAD3*×Aβ interaction using genetically predicted expression data as well as a (C) *TRIM65*×Aβ×sex interaction using RNAseq and (D) *TRIM65*×Aβ×sex interaction using genetically predicted expression data. The mean ± 1 standard deviation (SD) Aβ burden is given such that blue represents the mean Aβ burden, red is 1SD above the mean, and green is 1SD below the mean.

### Overlap with bulk brain RNAseq (ROSMAP)

Finally, we examined the brain relevance of our whole blood gene candidates from the gene×Aβ-PET and gene×Aβ-PET×sex models. To do this, we used bulk RNAseq data from dorsolateral prefrontal cortex (DLPFC) from 890 individuals enrolled in the Religious Orders Study and Rush Memory and Aging Project (ROSMAP)^52, 53^. To represent Aβ-PET, we used immunohistochemically derived measures of Aβ load at autopsy. For pTau217, we ran models with both immunohistochemically and silver-staining derived measures of tangle density and neurofibrillary tangle (NFT) burden as outcomes. Of the 2,275 unique genes represented in our primary whole blood analyses, 1,545 were available to test in ROSMAP after quality control.

No genes showed a significant three-way interaction between gene expression, sex, and Aβ burden on NFT burden or tangle density in brain tissue. In gene×Aβ models where NFT burden was the outcome, we found that 88 genes from brain tissue reached significance (p_FDR_ < 0.05), with 29 showing the same direction of effect as in whole blood. When using tangle density asbthe outcome in models, 147 genes from brain tissue reached significance in gene×Aβ models, with 69 showing the same direction of effect as in whole blood (**Supplementary Table 10**). 18 genes were common across both tau outcomes.

Significant genes overlapping between brain tissue and whole blood implicated processes such as ubiquitination (*KLHL2*)^54^, transcriptional regulation (*ZNF143*)^55^, and RNA splicing (*CASC3*)^56^. One X-linked gene, *KDM6A*, showed a significant interaction with brain Aβ burden on NFT burden (β=0.10(0.03), p_FDR_=0.03) and tangle density (β=0.10(0.03), p_FDR_=0.03) in these analyses (**Figure 5**). The most significant of the genes showing overlap between whole blood in A4/LEARN and DLPFC in ROSMAP were *KLHL2* (NFT burden, β=0.13(0.03), p_FDR_=3.34×10^−4^) and *C5orf22* (tangle density, β=-0.17(0.03), p_FDR_=3.19×10^−4^).

**Figure 5.**
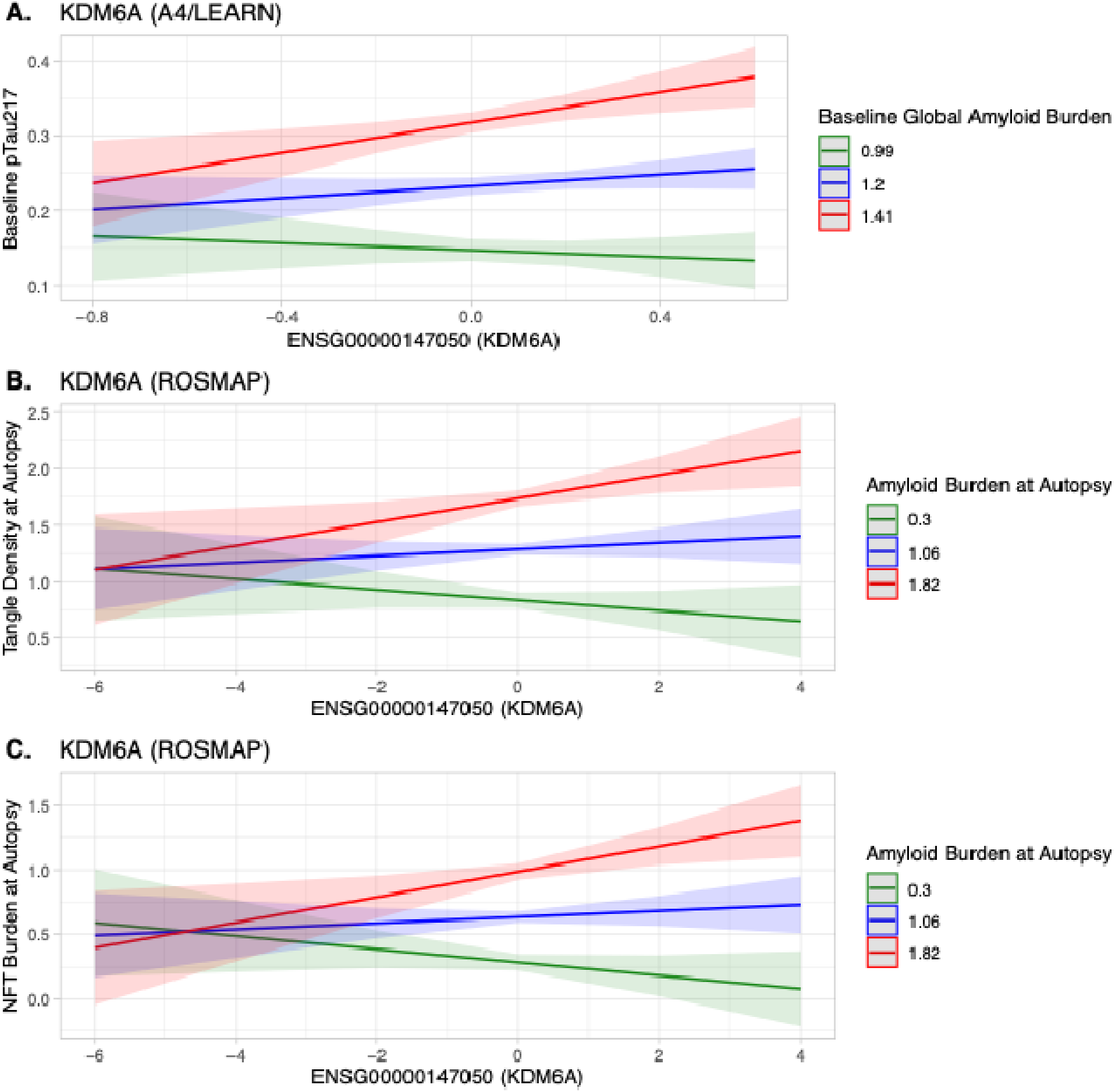
Model estimates predicting tau burden from an (A) *KDM6A*\*Aβ interaction in A4/LEARN and (B) *KDM6A*\*Aβ interaction in ROSMAP (tangle density) and (C) NFT burden. Models in A4 were adjusted for age, sex, BMI, and cohort; models in ROSMAP were adjusted for age of death, sex, and post-mortem interval. The mean ± 1 standard deviation (SD) Aβ burden is given such that blue represents the mean Aβ burden, red is 1SD above the mean, and green is 1SD below the mean.

Overrepresentation analyses via the Gene Ontology database (PANTHER Overrepresentation Test, https://functionome.geneontology.org/)^57^ on these overlapping genes provided GO:BP terms such as “cell cycle process”, “cellular localization”,” “embryonic viscerocranium morphogenesis”, and “fatty acid oxidation,” though none were statistically significant after correcting for multiple comparisons (**Supplementary Table 11**).

## Discussion

In this study of 724 cognitively unimpaired older adults, we examined whole blood transcriptomic correlates of plasma pTau217, a biomarker that has emerged as one of the most sensitive indicators of preclinical AD. We found no direct gene associations with pTau217 itself. Instead, more than 1,500 genes showed associations moderated by Aβ burden supporting the hypothesis that elevation in plasma pTau217 reflects systemic responses to Aβ-related cellular stress and dysregulation early in the disease process. These included known AD risk genes such as *EPHA1* and *TREML2*^29–34^ as well as novel candidates linked to cell metabolism and transcriptional regulation. Roughly 63% of these genes exhibited protective effects such that higher gene expression was related to lower plasma pTau217 burden among individuals with elevated Aβ-PET, which may inform therapeutic discovery for AD. Using the MetaXcan method, we find evidence of over 40 gene associations that may be genetically as opposed to pathologically driven, highlighting potential avenues for early risk detection. Our study also identified 80 unique genes that were significant and showed consistent directionality in both whole blood RNAseq data from A4/LEARN and bulk DLPFC RNAseq from ROSMAP. These results highlight the brain relevance of our study and suggest that biological functions such as protein degradation, RNA splicing, cell cycle regulation, and lipid metabolism may be relevant for AD in both tissues.

No genes were directly moderated by sex in our study, though 772 genes demonstrated sex-moderated associations with plasma pTau217 in the presence of Aβ. Our gene×Aβ-PET×sex findings are consistent with previous studies by our group showing that sex differences in AD risk (tau-PET accumulation and cognitive decline) are primarily driven by individuals with elevated Aβ^19–21^. Significant genes moderated by both Aβ and sex were implicated in cellular metabolism, immunity, and RNA processing. In these three-way interaction models, the most significant genes were *AMIGO2* and *CERKL*, which showed opposite directions of association by sex. These genes have been implicated in axonal guidance and mitochondrial function^39, 40^, respectively. Together, these findings suggest that Aβ burden may provide a critical biological context in which sex-specific transcriptional associations with plasma pTau217 become apparent.

A key innovation of this study was the systematic focus on the X chromosome, which has been understudied in aging and AD. We identified over 100 X-linked genes that significantly interacted with Aβ or both Aβ and sex on plasma pTau217, supporting a role for the X chromosome in shaping the early stages of AD progression. The X-linked genes moderated by Aβ-PET included *KLHL13*^*58*^ and *AIFM1*^*59*^, implicating functions such as ubiquitination, apoptosis, and mitochondrial function in the appearance of plasma pTau217. Among the significant X-linked genes found in our gene×Aβ-PET×sex models, hits such as *HK2P1* and *WWC3* converged on cellular signaling and glycolysis^43^, suggesting possible molecular pathways through which sex may shape vulnerability to tau phosphorylation. Notably, *KDM6A*, emerged consistently across blood and brain gene×Aβ-PET analyses. *KDM6A* has been previously implicated in studies of aging and AD, and it has been linked to resilience in animal studies and slower cognitive decline in humans^36, 60^. By contrast, our results suggest greater expression of *KDM6A* is associated with higher levels of both plasma pTau217 and tau burden at autopsy among individuals with elevated Aβ. This work warrants further investigation and continues to reinforce the importance of X chromosome exploration in AD research.

Genetically predicted expression (via MetaXcan)^47^ of over 40 genes showed significant and consistent interactions with Aβ-PET and sex on plasma pTau217, suggesting that some of these genes in whole blood are not simply demonstrating modification by AD pathology, and thus may be useful for early risk stratification and intervention. For instance, two genes of interest were *SMAD3* and *TRIM65*. Both are implicated in inflammation and immunity, which has been strongly linked to AD pathophysiology. *SMAD3* functions in the TGF-β pathways and plays a role in microglial and astrocytic function, particularly in the activation of Aβ uptake and clearance from the brain^49, 50^. A recent study found that higher blood *SMAD3* was associated with lower brain Aβ, consistent with our results^61^. *TRIM65* was significantly moderated by both Aβ-PET and sex in our findings; *TRIM65* negatively regulates the *NLRP3* inflammasome^51^; activation of *NLRP3* and neuroinflammation has been observed in animal models and individuals with AD pathology^62^. Therapies targeting the *NLRP3* inflammasome pathway have reduced Aβ deposition and memory impairment in AD mouse models^62^, supporting this pathway as a possible sex-specific therapeutic target for AD.

Our study leverages the advantages of transcriptomic data derived from whole blood. Blood draws are both minimally invasive and scalable, providing an opportunity to assess systemic biological processes that may relate to AD dementia and brain aging in more individuals. Despite this advantage, several limitations warrant discussion. Whole blood transcriptomic data may not fully reflect brain-specific processes, although they likely capture systemic immune and metabolic states relevant to AD. Although there were genes that were significant in both whole blood and brain-derived data, concordance was partial (5%), reflecting the deviation between whole blood pTau217 and neuropathological measures of Aβ load, tau neurofibrillary tangles and tangle density. This underscores both the complementary and distinct biology captured by peripheral versus central tissues and highlights the need for future studies that integrate blood- and brain-based transcriptomics with other multi-omic layers in longitudinal cohorts. Second, sample sizes remain modest for our interaction models, and larger independent datasets will be needed to confirm these effects. Third, our analyses were cross-sectional and cannot establish causation. Finally, our analyses were run on a cohort that is low in racial and ethnic diversity, leaving an open question as to how these findings might replicate in a more population-representative sample.

Despite these caveats, the present study provides novel evidence that peripheral transcriptomic signatures, moderated by Aβ and sex, are associated with plasma pTau217 levels. By implicating pathways in cell metabolism, transport, immunity, and gene regulation among others, this work advances a more nuanced view of plasma pTau217 as a biomarker and highlights promising new avenues for mechanistic discovery and sex-informed risk stratification in AD.

## Online Methods

### Participants

We analyzed data from 724 cognitively unimpaired participants drawn from the Anti-Amyloid Treatment in Asymptomatic Alzheimer’s (A4) clinical trial (n = 445) and the adjoining Longitudinal Evaluation of Amyloid Risk and Neurodegeneration (LEARN) observational study (n = 279)^63, 64^. Individuals with elevated Aβ-PET at screening (Aβ-PET standardized uptake value ratio, SUVR ≥ 1.15 and qualitative visual read) and who met all inclusion criteria were enrolled in the A4 trial; individuals with an Aβ-PET SUVR between 1.10 and 1.15 were also considered to be positive for enrollment with a positive visual read^28^. Individuals who were otherwise eligible for A4 but did not show elevated Aβ-PET were eligible to enroll in the LEARN study. All participants in A4 and LEARN underwent the same cognitive and functional assessments^16^. All data for the present study come from the pre-randomization screening visit for the A4 clinical trial. Eligibility for the current, primary analysis required availability of a single ^18^F-florbetapir Aβ-PET scan, plasma pTau217 measures, whole blood RNAseq data, and *APOE* genotype.

The A4 study protocol was approved by institutional review boards (IRBs) at each study site, with all participants providing written informed consent. This work was conducted in accordance with ethical guidelines of the Mass General Brigham (MGB) Human Research Committee. Deidentified data are publicly available at https://www.a4studydata.org and via www.synapse.org. RNA sequencing (RNAseq) data can be found at https://atri-a4.atrihub.org/docs/mgh_rnaseq.

### Whole blood RNAseq

At trial screening, whole blood (2.5 mL) was collected in PAXgene tubes, frozen on site, shipped on dry ice, and stored at −80°C until processing. RNA extraction was performed using the QIASymphony RNA Kit (QIAGEN, 931636), followed by depletion of ribosomal RNA and hemoglobin with the NEBNext Globin and rRNA Depletion Kit (New England BioLabs, E7750). Libraries were prepared using the NEBNext Ultra Directional Library Prep Kit (New England BioLabs, E7420) and sequenced on an Illumina NovaSeq 6000 using 151 bp paired end reads, targeting an average depth of 60 million reads per sample. All sequencing was carried out at the VANTAGE core at Vanderbilt University Medical Center (Nashville, TN, USA). Quality control and preprocessing were conducted by the Vanderbilt Memory and Alzheimer’s Center using established bulk RNAseq pipelines^23^. Briefly, gene-level counts were quantile normalized and adjusted for technical variation (e.g., batch effects) according to these standards. Counts for X-linked genes were normalized and adjusted separately from autosomal gene counts.

### Plasma pTau217

Screening samples of plasma pTau217 were processed by Eli Lilly and Company using an automated electrochemiluminescent immunoassay (Tecan Fluent workstation for preparation, MSD Sector S Imager 600MM for detection)^10^. Data are represented as pg/mL.

### Aβ-PET imaging

All participants in this study underwent Aβ-PET imaging with ^18^F-Florbetapir (10 mCi), acquired 50–70 minutes post-injection^65^. Aβ burden was quantified as a mean cortical SUVR using the whole cerebellum as the reference region. Aβ status was determined using a primary threshold of SUVR ≥ 1.15 to define Aβ positivity^64^.

### Primary statistical analyses

Linear regression models adjusting for age, body mass index (BMI – to account for blood volume), and cohort (i.e., A4, LEARN) examined associations between the following terms and pTau217 (pg/mL): gene, gene×Aβ-PET, gene×*APOE*ε4 status, gene×sex, and gene×Aβ-PET×sex. Positive *APOE*ε4 status was defined as individuals carrying at least one ε4 allele. Results for each model were corrected for the number of tested autosomal and X-linked genes (n_total_=20,621; 19,573 autosomal, 1,048 X-linked) using the false discovery rate (FDR) method. Sensitivity analyses included subgroup analysis (A4 vs LEARN). Stratified models (i.e., sex-stratified, *APOE*ε4-stratified) were examined *post hoc. Post hoc* analyses on individual genes were not corrected for multiple comparisons unless otherwise stated. Additional exploratory analyses are described below.

### Gene set enrichment analyses

Pre-ranked gene set enrichment analyses based on beta coefficients from each regression model were performed using the R package *fgsea* (v1.26.0) and Gene Ontology: Biological Process terms (GO:BP, downloaded October 31, 2025^37^) containing a minimum of 15 genes and a maximum of 500 genes. Significance for all analyses was set *a priori* at FDR-corrected p < 0.05. All significant transcriptomic results are available in **Supplementary Materials**.

### Genetically predicted gene expression (PrediXcan)

The PrediXcan method (https://github.com/hakyimlab/PrediXcan) was used to predict genetically regulated gene expression (i.e., the proportion of gene expression only attributed to genetic variation and not environment or phenotype) from genotype data in our sample. Genotype data was derived from blood using the Illumina Global Screening Array (build 37) and followed a previously reported imputation and quality control pipeline^66^. Briefly, variants with genotyping efficiency < 95% and minor allele frequency < 1% were excluded. Samples with call rates < 99%, inconsistencies between reported and genetic sex, or high relatedness (pi-hat > 0.25) were removed. Genotype data was lifted-over to build 38 and restricted to Non-Hispanic White individuals prior to imputation on the TOPMed Imputation Server (https://imputation.biodatacatalyst.nhlbi.nih.gov). PrediXcan models were built with [whole blood] gene expression data from GTEx (version 8, build 38; NIH database of Genotypes and Phenotypes (dbGaP) accession number: phs000424.GTEx.v8.p2, downloaded 11/13/2019) following a previously described method utilizing an elastic-net supervised machine learning model (α = 0.5, equal weights for LASSO and ridge regression) and applied to quality controlled genotype data^47^. Predicted gene expression from whole blood was available for 637 participants from our original sample. These data were used in the same models specified in ***Statistical Analyses***.

### Dorsolateral prefrontal cortex RNAseq

Gene expression data derived from dorsolateral prefrontal cortex (DLPFC) were also drawn from the Religious Orders Study and the Rush Memory and Aging Project (ROSMAP; n=890, **Supplementary Table 12**)^52, 53^ to identify overlapping signals between blood and brain tissue. ROSMAP enrolls older adults without known dementia who agree to annual clinical evaluations and brain donation at death. All participants provided written informed and repository consents, and an Anatomic Gift Act. Procedures for both studies were approved by an Institutional Review Board of Rush University Medical Center, with secondary analyses of existing data approved by the Vanderbilt University Medical Center IRB. DLPFC tissue was acquired at autopsy. Full details on RNA extraction, library preparation, and RNA sequencing have been described previously. Sequence alignment and quality control of data followed the same protocol as described for A4 RNAseq. ROSMAP data are available through the Accelerating Medicines Partnership – Alzheimer’s Disease (AMP-AD) Knowledge Portal (https://adknowledgeportal.synapse.org/Explore/Studies/DetailsPage/StudyDetails?Study=syn3219045) and the Rush Alzheimer’s Disease Center Resource Sharing Hub (https://www.radc.rush.edu/).

### ROSMAP brain neuropathological measures

Measures of neurofibrillary tangle burden, PHFtau tangle density, and β-amyloid load at autopsy were used in analyses using ROSMAP DLPFC RNAseq in proxy of plasma pTau217 and Aβ-PET, respectively. Neurofibrillary tangle burden was determined via examination of silver-stained slides from 5 brain regions (midfrontal cortex, midtemporal cortex, inferior parietal cortex, entorhinal cortex, and hippocampus). PHFtau tangle density was obtained via immunohistochemistry using an anti-AT8 antibody from 8 brain regions (angular gyrus, anterior cingulate, calcarine cortex, entorhinal cortex, hippocampus, inferior temporal cortex, midfrontal gyrus, and superior frontal cortex) and square-root transformed prior to analysis^67^. Aβ load was obtained from the same 8 brain regions as tau via immunohistochemistry using 3 monoclonal antibodies against Aβ (4G8, 6F/3D, and 10D5)^67^ over the past 3+ decades. Additional information on methods and quantification can be found on the Rush Alzheimer’s Disease Center Resource Sharing Hub (https://www.radc.rush.edu/).

## Data Availability

A4/LEARN data are publicly available at https://www.a4studydata.org and via www.synapse.org. RNA sequencing (RNAseq) data can be found at https://atri-a4.atrihub.org/docs/mgh_rnaseq. ROSMAP data are available through the Accelerating Medicines Partnership – Alzheimer’s Disease (AMP-AD) Knowledge portal (https://adknowledgeportal.synapse.org/) under accession code syn3219045. Data are available under controlled use conditions due to privacy laws. Access can be obtained by requesting a data use agreement on the AMP-AD portal or from the Rush Alzheimer’s Disease Center Resource Sharing Hub (https://www.radc.rush.edu/requests.htm).

## Article Information

### Conflict of Interest

RAS has served as a consultant for AbbVie, AC Immune, Acumen, Alector, Apellis, Biohaven, Bristol Myers Squibb, Ionis, Janssen, Oligomerix, Prothena, Roche, and Vaxxinity over the past 3 years. She has received research funding from Eisai and Eli Lilly for public-private partnership clinical trials and receives research grant funding from the National Institute on Aging/National Institutes of Health, GHR Foundation, and the Alzheimer’s Association. Her spouse, K. Johnson, reports consulting fees from Novartis, Merck, and Janssen. No other authors have disclosures relevant to this manuscript.

### Author Contributions

*Concept and design:* Seto, Buckley

*Acquisition, analysis, or interpretation of data:* All authors.

*Drafting of the manuscript:* Seto, Buckley

*Critical revision of the manuscript for important intellectual content:* All authors.

*Statistical analysis:* Seto, Klinger *Obtained funding:* Buckley, Sperling *Supervision:* Buckley

### Funding

This work was funded by the United States National Institutes of Health, DP2AG082342 [R.F.B.]), R01AG073439 [L.D.], R01AG063689 [R.A.S. and others], and U19AG010483 [R.A.S. and others]. The A4 Study is funded by NIH grants, Eli Lilly and Co, and several philanthropic organizations. ROS/MAP is supported by the following grants from the National Institute on Aging: P30AG10161, R01AG15819, R01AG17917, R01AG30146, R01AG36042, RC2AG036547, R01AG36836, R01AG48015, RF1AG57473, U01AG32984, U01AG46152, U01AG46161, U01AG61356 as well as the Illinois Department of Public Health, and the Translational Genomics Research Institute.

## Acknowledgements

We thank the participants and study staff of the A4 Study as well as ROSMAP. The Genotype-Tissue Expression (GTEx) Project was supported by the Common Fund of the Office of the Director of the National Institutes of Health, and by NCI, NHGRI, NHLBI, NIDA, NIMH, and NINDS.

## Notes

### Summary of Updates

Additional analyses using genetically predicted gene expression via MetaXcan, sensitivity analyses stratifying by cohort, and post hoc analyses with bulk brain RNAseq.

